# Plasma Amino Acid Responses to an Oral Glucose Challenge Relate More Strongly to Body Adiposity Than to Insulin Resistance

**DOI:** 10.64898/2026.06.14.732197

**Authors:** Eduardo D. S. Freitas, Kailin A. Johnson, Matthew Buras, Lori R. Roust, Eleanna De Filippis, Brooke B. Brown, Christos S. Katsanos

**Author notes:** Correspondence: Christos S. Katsanos, PhD, Health Futures Center, Room 331C, 6161 E. Mayo Blvd, Phoenix, AZ 85054.

## Abstract

The coexistence of obesity and insulin resistance is associated with elevated plasma amino acid concentrations. However, it remains unclear whether adiposity or insulin resistance is the stronger determinant of plasma amino acid dysregulation in this setting. Twenty-two adults (10 women, 12 men) spanning a broad range of body mass index (BMI) and insulin resistance underwent a 75-g oral glucose tolerance test (OGTT) after an overnight fast. Plasma glucose, insulin, and amino acid concentrations were measured serially, and insulin resistance/sensitivity was estimated from OGTT-derived glucose and insulin responses, using the homeostasis model assessment of insulin resistance (HOMA-IR) and the Matsuda insulin sensitivity index (Matsuda-ISI). Principal component analysis (PCA) of fasting plasma amino acid concentrations showed no clear separation by obesity or insulin resistance classifications. In contrast, PCA of OGTT-stimulated plasma amino acid concentrations revealed clearer clustering by BMI, fat mass, and waist circumference, whereas separation by HOMA-IR and Matsuda-ISI was less distinct. Importantly, regression analyses showed that BMI, fat mass, and waist circumference were significant predictors of OGTT-stimulated, but not fasting, amino acid responses, with waist circumference accounting for the greatest proportion of the variance in branched-chain amino acid responses during the OGTT (R^2^ = 0.54). In conclusion, measures of adiposity, particularly total fat mass and waist circumference, accounted for a greater proportion of the variance in plasma amino acid responses under physiologically stimulated conditions than indices of insulin resistance. These findings support the view that plasma amino acid concentrations reflect adiposity-related metabolic alterations more strongly than insulin resistance.

## INTRODUCTION

Elevated circulating amino acid concentrations represent one of the most reproducible metabolic signatures of human obesity and insulin resistance, with the most consistent increases observed in branched-chain amino acids (BCAA) and aromatic amino acids (1–3). Alterations in plasma amino acid profiles have largely been attributed to the presence of insulin resistance in these individuals (4, 5). However, because insulin resistance commonly coexists with increased adiposity in these individuals (6, 7), existing studies have not been able to disentangle the relative contributions of adiposity and insulin resistance per se to elevated circulating amino acid concentrations. Notably, elevated BCAA concentrations have themselves been implicated in the development of insulin resistance (8).

Excess adiposity may be a primary driver of elevated circulating amino acids in obesity through both quantitative and qualitative effects of expanded fat mass. For example, adipose tissue possesses a measurable capacity to catabolize circulating BCAA, thereby can contribute to systemic BCAA homeostasis (9). At the same time, adipose tissue expansion in obesity is accompanied by downregulation of genes encoding key components of the BCAA catabolic pathway (10). Therefore, reduced BCAA metabolic capacity with expanded adiposity may contribute directly to higher plasma amino acid concentrations, even in the absence of systemic insulin resistance. On the other hand, insulin is a potent regulator of amino acid metabolism, exerting inhibitory control over whole-body, and particularly skeletal muscle, protein breakdown (11). In insulin-resistant states, this suppression of proteolysis is blunted (12), which can consequently lead to increased amino acid release into the circulation. Thus, insulin resistance may elevate plasma amino acids primarily through increased plasma appearance secondary to impaired insulin-mediated control of protein breakdown.

Disentangling the relative contributions of adiposity versus insulin resistance to plasma amino acid metabolism is of both physiological and clinical significance. A major limitation of the existing literature is that the evidence linking plasma amino acid concentrations with obesity and insulin resistance has been obtained under basal, fasting conditions. The fed state may be more physiologically informative than the fasting state for evaluating plasma amino-acid responses. This is because the fed state places greater demands on nutrient metabolism (13), and, thus, postprandial plasma amino acid responses can provide a more sensitive and informative window into whether excess adiposity versus insulin resistance is the dominant determinant of circulating amino acid concentrations. Consistent with this concept, metabolic inflexibility in individuals with obesity and insulin resistance is most apparent during the postprandial period, including more prolonged elevations in plasma amino acids (14).

The present study was therefore designed to determine whether adiposity-related measures, rather than insulin resistance, are more closely related to plasma amino acid concentrations under basal and nutrient-stimulated conditions, with a standard oral glucose tolerance test (OGTT) serving as the dietary challenge. If adiposity-associated parameters account for a larger share of the observed variability in plasma amino acid concentrations than indices of insulin resistance, then observed disruption in plasma amino acid homeostasis may be viewed as a downstream metabolic manifestation more closely linked to adiposity.

## METHODS

### Study Participants

The study cohort was intentionally designed to include participants with a wide range of body mass index (BMI) values in order to capture broad variation in adiposity and insulin resistance, and given the well-established association between BMI, adiposity, and insulin resistance (6, 7). Participants had no evidence of acute illness and no history of, or evidence on screening for, diabetes or heart, liver, kidney, or gastrointestinal disease. Participants were also excluded if they were taking any medications or over-the-counter supplements. All study participants were physically inactive according to current guidelines, defined as engaging in regular physical activity on no more than two days per week (15, 16). None of the participants were actively attempting weight loss. Before undergoing any study procedures, participants were informed of all experimental procedures and the risks associated with study participation, and written informed consent was obtained. The study was reviewed and approved by the Mayo Clinic Institutional Review Board.

### Overall Study Design

All data were collected during a morning study visit at the Ambulatory Infusion Center at Mayo Clinic in Arizona after an overnight fast of 10–12 hours. The study visit included body composition assessment by bioelectrical impedance analysis (BIA 310e, Biodynamics Corp.), blood collection for measurement of fasting biochemical parameters, and completion of an OGTT (17). Although the OGTT does not provide a conventional meal-related stimulus, it constitutes a well-defined physiological stimulus that challenges insulin-regulated substrate, including amino acid, metabolism, thereby revealing differences in endogenous amino acid regulation under standardized conditions. For the OGTT, an intravenous catheter was placed in an antecubital vein for serial blood sampling while participants rested in a medical recliner chair. After approximately 15 min of rest, a baseline blood sample was obtained, followed immediately by ingestion of 75 g of glucose dissolved in 250 mL of water. Additional blood samples were collected at 30, 60, 90, and 120 min after glucose ingestion. Blood samples were centrifuged at 4 °C for ∼ 10 min at 3000 x g to separate plasma, and plasma aliquots were stored in plastic microcentrifuge tubes at -80 °C until analyses.

### Determination of Glucose and Insulin Concentrations and Calculation of Insulin Resistance/Sensitivity Indices

Glucose concentrations (mg/dL) were measured using an automated glucose analyzer (YSI 2300, Yellow Springs, OH). Insulin concentrations (mIU/mL) were measured by enzyme-linked immunosorbent assay (80-INSHU-E10.1, ALPCO Diagnostics). Fasting glucose and insulin values were used to calculate the Homeostasis Model Assessment of Insulin Resistance (HOMA-IR) (18). Glucose and insulin values obtained during the OGTT were used to calculate the Matsuda Insulin Sensitivity Index (Matsuda-ISI), an estimate of whole-body insulin sensitivity (17).

### Determination of Amino Acids Concentrations

Plasma amino acid concentrations were measured by LC-MS/MS at the Mayo Clinic Metabolomics Core following derivatization with 6-aminoquinolyl-N-hydroxysuccinimidyl carbamate using the AccQ-Fluor™ kit (Waters Corporation), as previously described (19). Amino acids measured included branched-chain amino acids (BCAA; leucine, isoleucine, and valine), essential amino acids (EAA; histidine, isoleucine, leucine, lysine, methionine, phenylalanine, threonine, tryptophan, and valine in addition to BCAA), and non-essential amino acids (NEAA; alanine, arginine, asparagine, aspartic acid, glutamic acid, glutamine, glycine, serine, and tyrosine).

### Statistical Analyses

Unpaired t-tests were used to compare group differences in biochemical and anthropometric characteristics at basal. Principal component analyses (PCA) were performed on plasma amino acid concentrations to examine underlying patterns in amino acid profiles. For visualization purposes, participants were stratified into groups of equal size using mean splits of obesity-related variables (BMI, fat mass, body fat percentage, and waist circumference) and insulin resistance/sensitivity indices (HOMA-IR and Matsuda-ISI). The overall significance of the PCA and the significance of individual principal components were evaluated in accordance with methods described by Camargo (20) and Vieira (21). Euclidean distances between clusters were calculated as the distances between cluster centroids based on PCA scores for the plotted principal components. Multiple linear regression analyses were performed to determine the extent to which adiposity-related and insulin resistance-related parameters explained variability in plasma amino acid concentrations in the fasting and OGTT-stimulated states. Statistical significance was set at *P* ≤ 0.05. Data are presented as means and ranges. Statistical analyses were performed using R Studio, version 4.3.2. (22).

## RESULTS

### Participant Characteristics

The study cohort (N = 22; 10 females, 12 males) exhibited broad range in body composition and insulin resistance measures, consistent with the study design, and summarized in **Table 1**. Consistent with prior reports (6, 7), higher adiposity (i.e., percent body fat) was associated with greater insulin resistance, as reflected by higher HOMA-IR in the fasting state and lower Matsuda-ISI in the OGTT-stimulated state (**Figure 1**). **Tables 2** and **3** show the plasma glucose and insulin responses as well as the plasma amino acid responses, respectively, under fasting conditions and during the OGTT-stimulated state.

**Figure 1.**
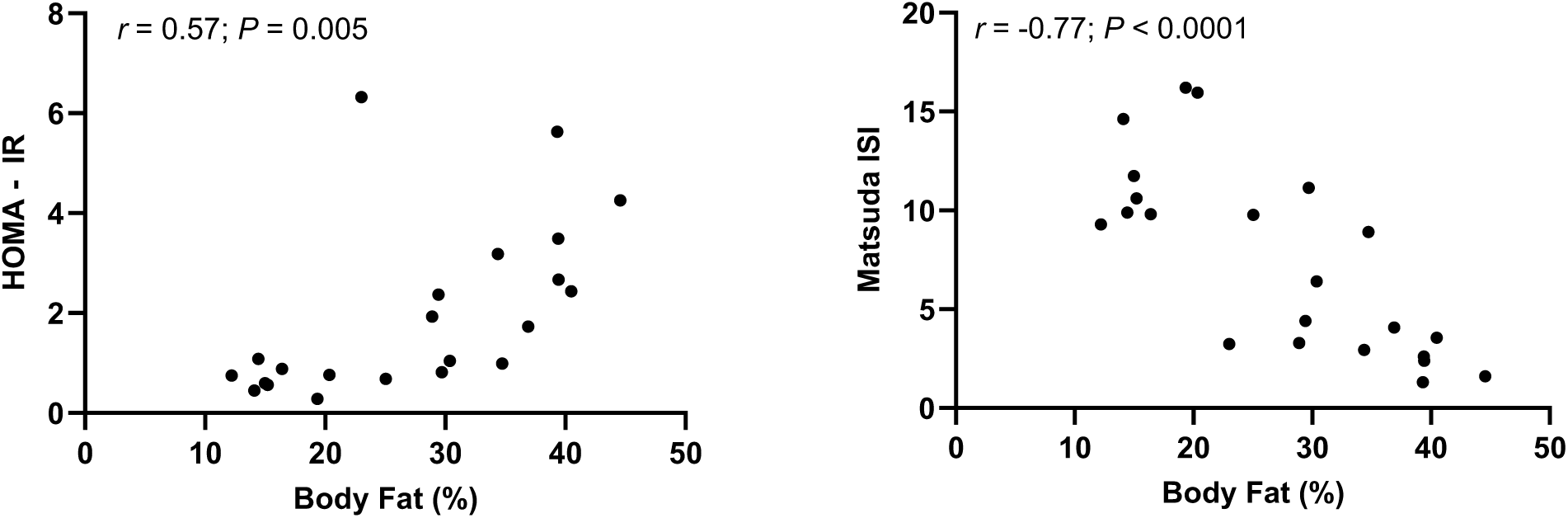
Associations between body fat percentage and indices of insulin resistance (HOMA-IR) and insulin sensitivity (Matsuda-ISI) across study participants. HOMA-IR, Homeostasis Model Assessment of Insulin Resistance; Matsuda-ISI, Matsuda Insulin Sensitivity Index. Pearson’s correlation coefficient (*r*) and corresponding *P* value are shown in each panel.

**Table 1.** Participant anthropometric and biochemical characteristics.

|  | Mean | Range |
| --- | --- | --- |
| Age (years) | 35.5 | (19.0 – 48.0) |
| Weight (kg) | 85.0 | (49.7 – 129.9) |
| Height (m) | 1.7 | (1.5 – 1.8) |
| BMI (kg/m<sup>2</sup>) | 28.8 | (19.4 – 41.0) |
| Waist circumference (cm) | 93.1 | (77.0 – 125.0) |
| Hip circumference (cm) | 107.1 | (82.5 – 140.3) |
| Waist-to-hip ratio (cm) | 0.8 | (0.7 – 1.1) |
| Fat-free mass (kg) | 58.3 | (34.6 – 80.4) |
| Fat mass (kg) | 24.6 | (7.7 – 57.9) |
| Body fat percentage (%) | 28.3 | (12.2 – 44.6) |
| VO<sub>2</sub>peak (ml/min) | 2103.9 | (916.2 – 3547.5) |
| VO<sub>2</sub>peak (ml/kgFFM/min) | 36.0 | (23.7 – 62.0) |
| Glucose (mg/dL) | 89.4 | (74.2 – 117.5) |
| Insulin (μIU/mL) | 8.7 | (1.4 – 27.8) |
| Matsuda-ISI | 7.3 | (1.3 – 16.2) |
| HOMA-IR | 1.97 | (0.28 – 6.32) |

Data are presented as mean and range. BMI, body mass index; FFM, fat-free mass; VO<sub>2</sub>peak, peak oxygen uptake; Matsuda-ISI, Matsuda insulin sensitivity index; HOMA-IR, homeostatic model assessment of insulin resistance.

**Table 2.** Plasma glucose and insulin concentrations during the oral glucose tolerance test.

|  | 0 min | 30 min | 60 min | 90 min | 120 min |
| --- | --- | --- | --- | --- | --- |
| Glucose (mg/dl) | 89.4 | 140.0 | 146.2 | 127.2 | 122.8 |
|  | (76.4 – 113.1) | (103.2 – 178.5) | (62.7 – 227.3) | (61.1 – 208.3) | (44.9 – 157.2) |
| Insulin (uIU/ml) | 9.3 | 53.9 | 62.5 | 69.4 | 56.9 |
|  | (1.4 – 27.8) | (14.9 – 160.6) | (16.9 – 246.6) | (5.8 – 265.4) | (5.6 – 164.2) |

Data are presented as mean and range

**Table 3.** Plasma amino acid concentrations during the oral glucose tolerance test.

|  | 0 min | 30 min | 60 min | 90 min | 120 min |
| --- | --- | --- | --- | --- | --- |
| Alanine | 370.9<br>(169.1 – 644.3) | 375.9<br>(218.3 – 654.2) | 375.2<br>(202.8 – 643.3) | 362.7<br>(219.2 – 591.8) | 342.5<br>(220.7 – 536.1) |
| Arginine | 71.5<br>(37.2 – 132.9) | 71.7<br>(35.3 – 115.4) | 68.2<br>(39.8 – 118.3) | 65.8<br>(36.3 – 102.8) | 65.8<br>(37.2 – 101.6) |
| Asparagine | 53.5<br>(23.7 – 87.5) | 53.7<br>(37.1 – 78.8) | 48.3<br>(33.3 – 72.6) | 44.2<br>(30.3 – 62.4) | 41.6<br>(20.1 – 59.9) |
| Aspartate | 12.7<br>(2.6 – 24.0) | 12.1<br>(3.8 – 22.6) | 10.3<br>(3.1 – 21.2) | 8.5<br>(3.2 – 14.8) | 7.1<br>(1.9 – 14.6) |
| Glutamine | 641.2<br>(318.3 – 1027.5) | 664.9<br>(393.1 – 1006.8) | 627.4<br>(349.8 – 941.6) | 597.4<br>(356.6 – 948.5) | 572.8<br>(335.2 – 863.4) |
| Glutamate | 51.9<br>(22.3 – 87.6) | 44.5<br>(20.3 – 84.5) | 38.4<br>(19.9 – 68.8) | 35.0<br>(15.8 – 68.2) | 32.3<br>(9.5 – 56.3) |
| Glycine | 218.2<br>(86.1 – 512.1) | 212.8<br>(130.4 – 455.5) | 191.9<br>(118.4 – 410.5) | 185.0<br>(118.5 – 348.1) | 171.0<br>(105.5 – 231.6) |
| Histidine | 76.4<br>(45.1 – 108.4) | 77.9<br>(58.3 – 100.9) | 72.5<br>(53.0 – 100.2) | 67.0<br>(51.9 – 83.1) | 65.0<br>(39.8 – 85.6) |
| Isoleucine | 58.7<br>(42.1 – 95.6) | 59.9<br>(41.4 – 108.5) | 50.4<br>(32.0 – 92.3) | 41.8<br>(22.8 – 81.3) | 36.3<br>(14.5 – 71.0) |
| Leucine | 113.4<br>(71.1 – 158.1) | 116.0<br>(82.7 – 175.7) | 99.0<br>(64.4 – 192.9) | 81.6<br>(55.7 – 130.9) | 70.6<br>(30.7 – 119.1) |
| Lysine | 181.8<br>(97.6 – 400.22) | 199.4<br>(74.5 – 483.6) | 179.7<br>(66.1 – 451.0) | 169.8<br>(77.7 – 491.9) | 152.0<br>(51.1 – 441.3) |
| Methionine | 23.8<br>(16.2 – 31.1) | 24.2<br>(16.0 – 32.5) | 22.1<br>(15.1 – 35.7) | 19.7<br>(12.4 – 25.5) | 17.7<br>(10.3 – 23.8) |
| Phenylalanine | 57.5<br>(41.3 – 75.3) | 58.7<br>(47.3 – 90.7) | 53.4<br>(36.0 – 85.5) | 48.2<br>(35.5 – 70.5) | 44.4<br>(28.6 – 64.5) |
| Serine | 102.7<br>(50.9 – 145.1) | 99.6<br>(62.3 – 134.0) | 90.2<br>(58.0 – 134.3) | 80.9<br>(48.6 – 106.6) | 72.9<br>(49.2 – 94.2) |
| Threonine | 153.9<br>(80.8 – 214.8) | 152.3<br>(82.1 – 205.7) | 137.9<br>(72.5 – 264.1) | 124.6<br>(66.3 – 184.12) | 115.3<br>(53.4 – 171.3) |
| Tryptophan | 50.6<br>(29.8 – 87.4) | 51.8<br>(33.1 – 87.4) | 49.7<br>(30.2 – 79.2) | 46.3<br>(28.2 – 74.1) | 43.7<br>(19.2 – 73.7) |
| Tyrosine | 57.5<br>(29.8 – 93.8) | 57.8<br>(29.9 – 101.1) | 52.8<br>(25.0 – 90.9) | 47.3<br>(23.1 – 81.1) | 42.8<br>(19.8 – 75.1) |
| Valine | 208.2 | 213.4 | 196.4 | 176.6 | 163.8 |
|  | (132.9 – 278.1) | (153.5 – 316.0) | (134.1 – 352.7) | (119.3 – 259.7) | (77.5 – 249.7) |

Data are presented as mean and range; values are in $\mu\text{mol/L}$ .

### Principal Component Analysis

Principal component analysis was used to reduce data dimensionality and identify underlying patterns in plasma amino acid responses that distinguish individuals with lower versus higher adiposity-related measures (BMI, fat mass, body fat percentage, and waist circumference) and indices of insulin resistance (HOMA-IR and Matsuda-ISI) stratifications. PCA of fasting plasma amino acid concentrations for individual amino acids was statistically significant (*P* < 0.05), with principal components 1 and 2 explaining 34.1% and 21.2% of the total variance, respectively. **Figure 2**, which displays the coordinates of each participant on principal components 1 and 2 derived from the PCA of fasting amino acid concentrations, shows no clear separation or clustering of participants according to adiposity or insulin resistance stratifications.

**Figure 2.**
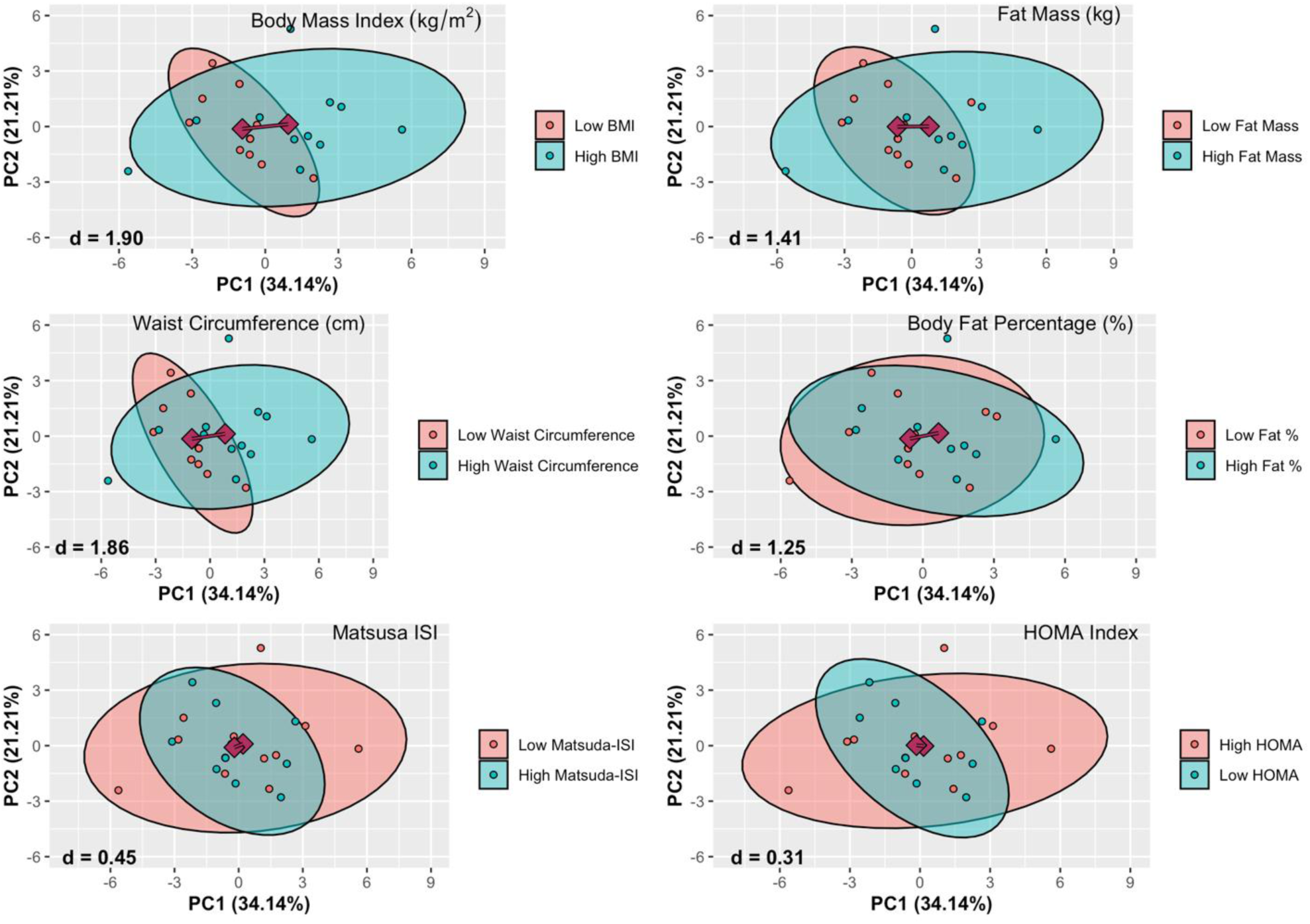
Principal component analysis of individual fasting plasma amino acid concentrations according to measures of obesity and insulin resistance. In each panel, the line represents the Euclidean distance between cluster centroids. BMI, body mass index; Matsuda-ISI, Matsuda insulin sensitivity index; HOMA-IR, homeostatic model assessment of insulin resistance.

We next performed PCA using plasma concentrations of individual amino acids averaged over the 2-h period following the OGTT. This analysis was statistically significant (*P* < 0.05), with principal components 1 and 2 accounting for 37.9% and 21.3% of the total variance, respectively. In contrast to the fasting state, PCA of plasma amino acid responses during the OGTT-stimulated state showed greater separation of participants in PCA space when stratified by BMI, fat mass, and waist circumference, but not body fat percentage, among the obesity measurements (**Figure 3**). By comparison, separation according to indices of insulin resistance (HOMA-IR and Matsuda-ISI) was less distinct, with substantial overlap between groups. Importantly, cluster separation, quantified as Euclidean distance between group centroids in PCA space, was consistently greater for OGTT-stimulated plasma amino acid responses (Figure 3) than for fasting plasma amino acids concentrations (Figure 2).

**Figure 3.**
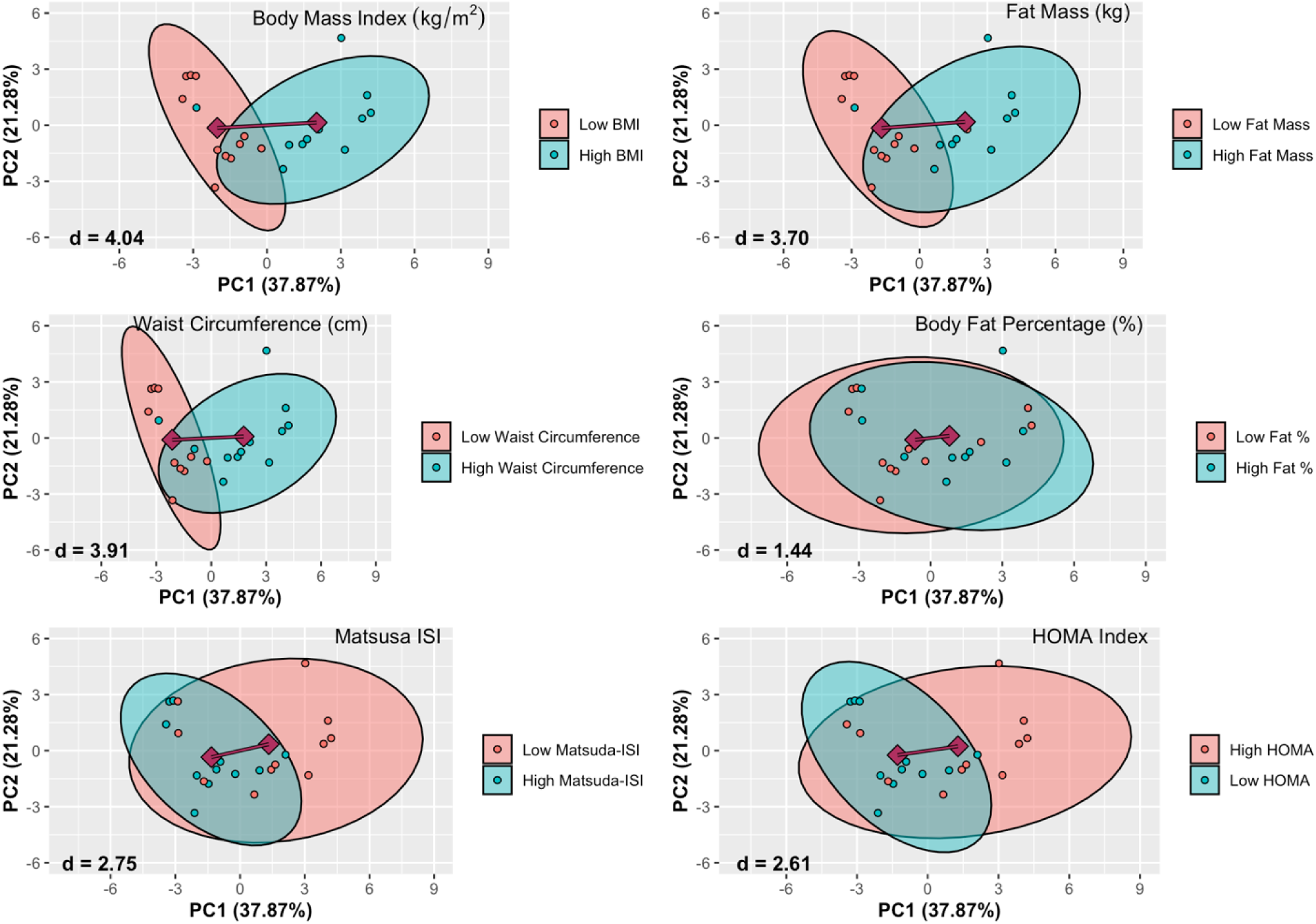
Principal component analysis of individual plasma amino acid responses during the OGTT-stimulated state according to measures of obesity and insulin resistance. In each panel, the line represents the Euclidean distance between cluster centroids. BMI, body mass index; Matsuda-ISI, Matsuda insulin sensitivity index; HOMA-IR, homeostatic model assessment of insulin resistance.

To assess the independent contributions of obesity and insulin resistance parameters to plasma amino acid concentrations, we performed multiple linear regression analyses separately for the fasting and OGTT-stimulated states, with obesity and insulin resistance measures entered as predictors and amino acid concentrations as the dependent variables. For the analyses, the plasma amino acids were grouped into total amino acids (TAA), EAA, BCAA, and NEAA. In the fasting state, waist circumference was the only significant predictor of plasma amino acid concentrations, and specifically BCAA concentrations (ß = 2.14, R^2^ = 0.186, *P* = 0.045), whereas no other significant predictors were identified for BCAA or any other plasma amino acid pool (all *P* > 0.05; data not shown). In contrast, in the OGTT-stimulated state, BMI, fat mass, and waist circumference were consistently significant predictors of TAA, BCAA, and EAA concentration responses (all *P* < 0.05; **Table 4**). By contrast, the insulin resistance/sensitivity indices HOMA-IR and Matsuda-ISI were not significant predictors of responses in any plasma amino acid pool (Table 4), and their inclusion in the models did not improve model fit or yield significant associations (*P* > 0.05).

**Table 4.**
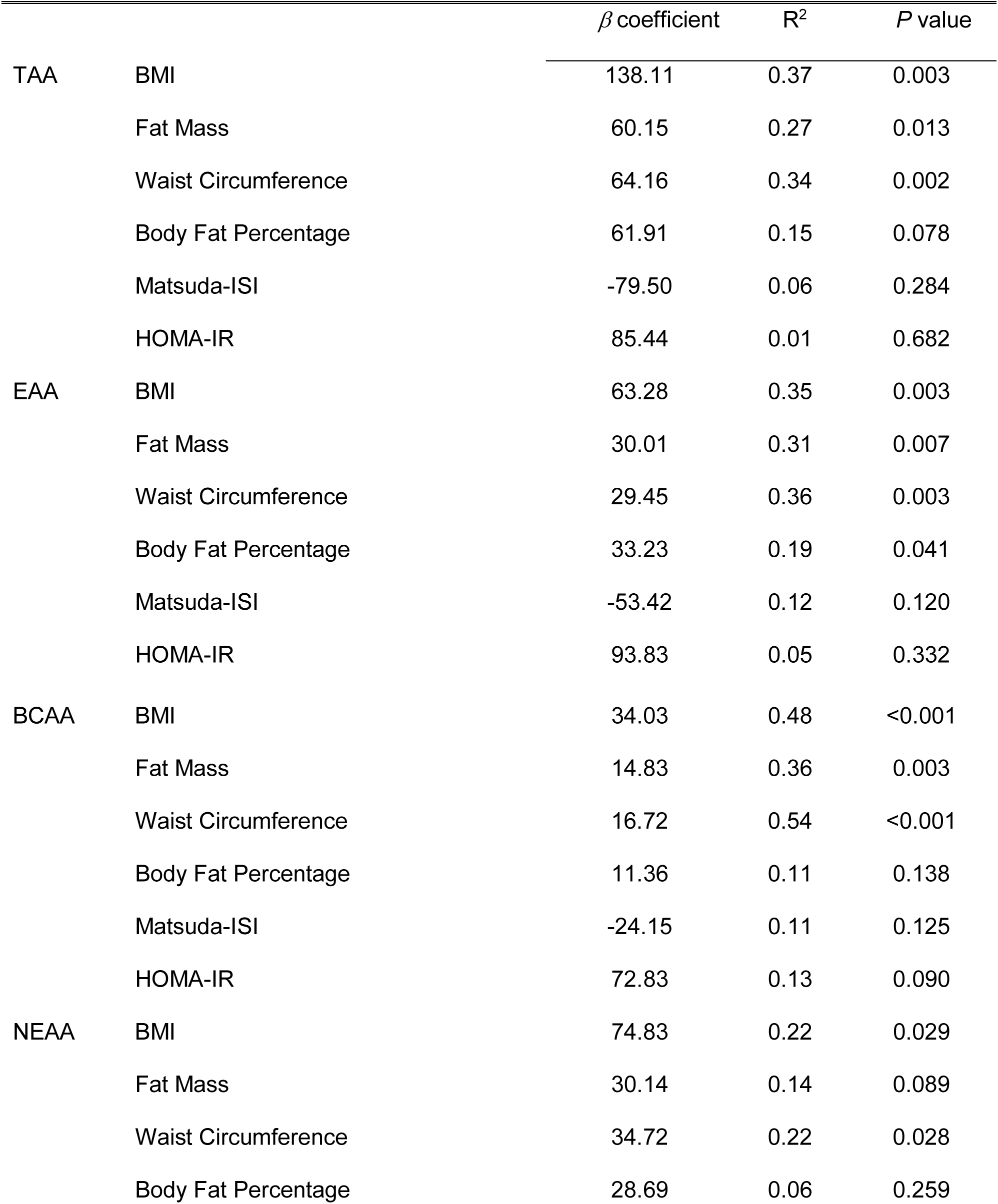
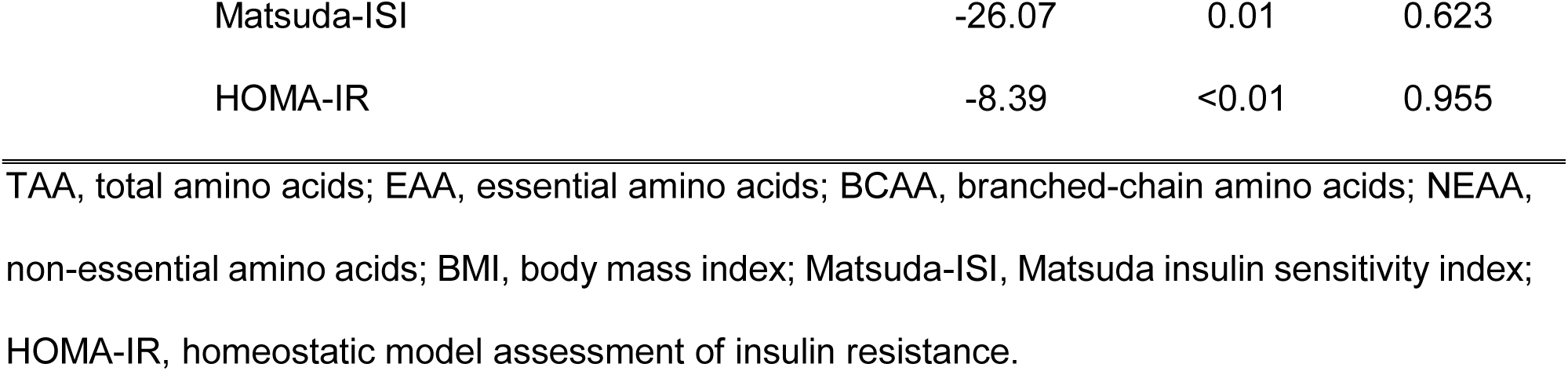
Linear regression models examining adiposity and insulin sensitivity parameters as predictors of plasma amino acid responses during the OGTT-stimulated state.

## DISCUSSION

Circulating amino acid concentrations, particularly BCAA, are consistently elevated in individuals with obesity and insulin resistance (1–3), yet the extent to which this phenotype is more closely related to excess adiposity or insulin resistance per se remains unresolved. This distinction is of physiological and clinical relevance, as elevated plasma amino acids have been proposed not only as biomarkers of metabolic dysfunction but also as potential contributors to the development of insulin resistance (8). Consistent with this concept, plasma amino acid concentrations have been shown to predict future insulin resistance, suggesting that altered amino acid metabolism may precede deterioration in glucose homeostasis (23). However, much of the evidence supporting these associations has been derived from fasting conditions, which may obscure physiological differences that manifest most clearly during nutrient-stimulated states, and given the evidence that metabolic inflexibility is often more evident under postprandial conditions (13, 14). The present study was therefore designed to compare the relative associations of obesity- and insulin resistance-related parameters with plasma amino acid concentrations under both fasting state and physiologically stimulated state induced by an OGTT.

The principal finding of this study is that plasma amino acid responses during the OGTT were more strongly associated with adiposity-related measures than with conventional indices of insulin resistance. In the fasting state, PCA revealed minimal separation of participants when stratified by either obesity-related measures or insulin resistance indices (Figure 2), indicating that fasting plasma amino acid profiles alone provide limited discriminatory insight across a broad spectrum of body composition and insulin sensitivity. In contrast, OGTT-stimulated plasma amino acid responses demonstrated greater separation in PCA space when participants were stratified by BMI, fat mass, and waist circumference, and whereas separation based on HOMA-IR or Matsuda-ISI remained modest with substantial overlap between groups (Figure 3). These visualization-based findings were reinforced statistically by regression analyses showing that adiposity-related variables, particularly fat mass and waist circumference, accounted for a greater proportion of variance in post-challenge amino acid concentrations than did indices of insulin resistance.

The observation that adiposity-related measures better accounted for variability in plasma amino acid responses during the OGTT than in the fasting state is consistent with the concept that metabolic inflexibility is more readily revealed under physiologically demanding than under basal, non-stimulated conditions (24–26). Glucose challenge imposes coordinated demands on nutrient metabolism, and the postprandial period therefore provides a more sensitive window into systemic metabolic regulation. Specifically, with respect to plasma amino acid responses, our data indicate that the elevations in plasma amino acid concentrations commonly observed in obesity are not accounted for by conventional indices of insulin resistance, as assessed by HOMA-IR and Matsuda-ISI, but instead are more closely related to measures of adiposity, particularly fat mass and waist circumference. In this context, our study reinforces previous evidence where elevation in plasma amino acid levels observed in humans with obesity was not associated with an impaired insulin-mediated amino acid flux (27). Notably, the obesity measures that best differentiated the stimulated amino acid response in the present study were BMI, fat mass, and waist circumference whereas body fat percentage was less informative. Therefore, the present data indicate that excess adiposity and its distribution are more closely linked to postprandial disturbances in plasma amino acid concentrations than are commonly used surrogate measures of insulin resistance. Among the obesity-related variables examined, waist circumference explained the greatest proportion of variance in BCAA concentrations during the OGTT, exceeding that of fat mass. This finding suggests that central fat distribution, followed by absolute adiposity, are more informative correlates of postprandial amino acid regulation than relative adiposity measures (i.e., percent body fat). Because a larger waist circumference largely reflects greater visceral adipose tissue accumulation, these findings extent prior evidence where visceral and intra-abdominal adipose tissue, more than subcutaneous adipose tissue, are associated with disturbed amino acid flux (28) and higher plasma amino acid concentrations (29). It is important to note that plasma amino acid concentrations reflect contributions from multiple tissues, including the liver, skeletal muscle, and adipose tissue. The present findings do not resolve the tissue-specific mechanisms underlying plasma amino acid regulation. However, given the associations of postprandial amino acid responses with adiposity-associated parameters, a biologically plausible framework for the observed associations centers on the role of adipose tissue in the observed plasma amino acid concentrations. It is known that adipose tissue contributes to systemic BCAA homeostasis (9), and obesity is associated with downregulation of genes involved in key steps of the BCAA oxidative pathway (10). Impaired adipose tissue BCAA catabolism has been proposed as a mechanism contributing to elevated circulating BCAA concentrations in obesity, and experimental disruption of these pathways increases plasma BCAA levels (30–32). While the present study did not directly assess tissue-specific amino acid flux, gene expression, or enzymatic activity, the convergence of our findings with existing literature supports the interpretation that excess adiposity, and particularly centralized fat accumulation, is closely linked to postprandial amino acid handling. Our findings therefore support a stronger link between adiposity-related measures and disturbances in plasma amino acid concentrations than between these disturbances and conventional indices of insulin resistance.

Consistent with our findings that plasma amino acid responses were not associated with insulin resistance, prior work has shown that acute elevations in circulating amino acids do not induce insulin resistance in healthy humans (33). Taken together, this body of evidence supports the concept that elevated plasma amino acids are not intrinsically detrimental to health. Indeed, higher habitual intakes of selected amino acids, including glutamic acid, leucine, and tyrosine, have been associated with favorable cardiovascular outcomes (34). Thus, elevated plasma amino acid concentrations in obesity may reflect downstream metabolic manifestations of excess adiposity occurring in concert with other adipose tissue-related disturbances, such as impaired suppression of lipolysis (35), rather than serving as independent or sufficient initiators of insulin resistance. Future studies combining dietary-stimulated amino acid responses with tracer or model-based approaches may help determine whether obesity-related differences in plasma amino acid responses reflect altered amino acid appearance, disposal, or oxidation, and illustrate how dynamic nutrient concentration data can be used to infer underlying substrate fluxes (36).

The finding that indices of insulin resistance did not independently predict plasma amino acid concentrations in either the fasting or OGTT-stimulated states, should be interpreted in the context of the measures used. For example, Matsuda-ISI provides a validated estimate of whole-body insulin sensitivity/resistance (17), but may not fully capture tissue-specific insulin actions that are most relevant to amino acid kinetics. Accordingly, the lack of strong associations with these indices does not imply that insulin resistance plays no role in amino acid metabolism, but rather that conventional insulin resistance metrics may be less informative determinants of interindividual variability in plasma amino acid concentrations than measures of adiposity in a physiological context. Moreover, if more precise measures of insulin resistance, such as the hyperinsulinemic-euglycemic clamp, were used, they assess insulin action under non-physiological conditions and do not capture the dynamic responses to a physiological nutrient challenge. We acknowledge that the present findings do not establish causality, but rather, demonstrate that obesity related variables account for a greater share of observed variability in plasma amino acid responses during an OGTT than do indices of insulin resistance in a heterogeneous cohort of adults without evidence of metabolic disease.

In conclusion, the present study demonstrates that plasma amino acid responses to a glucose challenge are more informative than fasting plasma amino acid concentrations and are more strongly associated with measures of adiposity, particularly fat mass and waist circumference, than with conventional indices of insulin resistance. These findings suggest that excess adiposity and its distribution are key correlates of postprandial amino acid dysregulation in humans. From a clinical perspective, our data suggest caution in using fasting plasma amino acid profiles alone as metabolic risk markers, and indicate that elevated plasma amino acid concentrations may be better interpreted as metabolic correlates of adiposity rather than insulin resistance.

## Author Contributions

EDSF and CSK conceived and designed the research; EDSF, KJ, LRR, EDF, BBB, and CSK performed the experiments; EDSF, KJ, MB, and CSK analyzed the data; EDSF and CSK interpreted the results of the experiments; EDSF and CSK prepared the figures; EDSF and CSK drafted the manuscript; EDSF, KJ, MB, LRR, EDF, BBB, and CSK edited and revised the manuscript; and EDSF, KJ, MB, LRR, EDF, BBB, and CSK approved the final version of the manuscript.

---

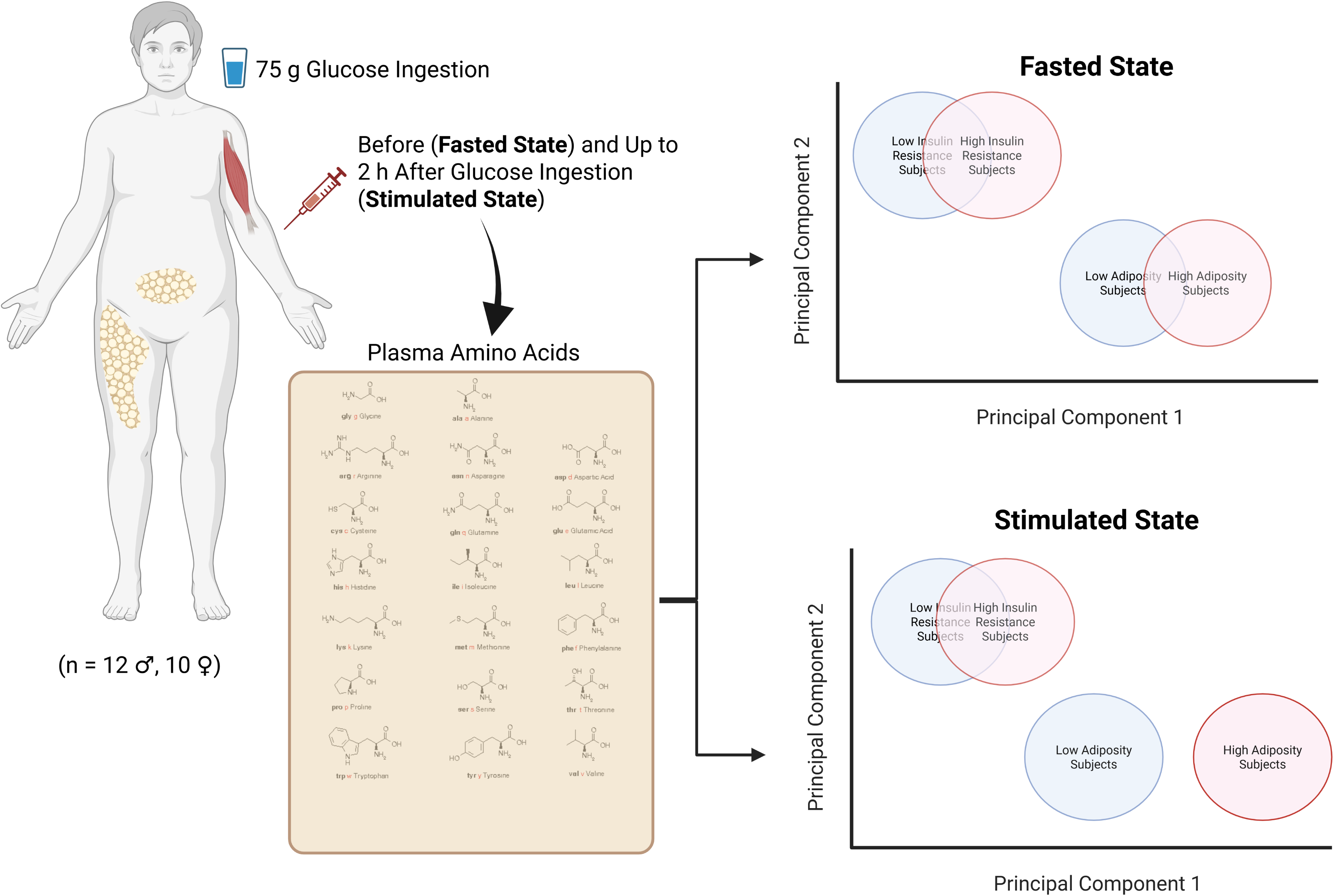

